# codonGPT: Reinforcement learning on a generative language model optimizes RNA sequences under biological constraints

**DOI:** 10.1101/2025.06.25.661500

**Authors:** Binita Rajbanshi, Anuj Guruacharya

## Abstract

Emerging generative models for biology focus on DNA, non-coding RNA, or proteins, ignoring information hidden in mRNA. Additionally, in protein engineering and mRNA therapeutics the design of mRNA sequences is still a challenge, lacking a clear framework. Here, we introduce and rigorously evaluate two novel methods: a foundational model for mRNA and a reinforcement learning mRNA design framework built on such a model. codonGPT is the first generative foundational language model trained directly on coding mRNA sequences. To solve the problem of synonymous constraints that are only unique to mRNA, we introduce a novel method of inference-time masking, along with house-keeping genes evaluation. For the first time, we also rigorously demonstrate, that for precise mRNA therapeutics design, reinforcement learning on such a model provides a clear framework for biological optimization. Our study introduces a novel foundational model for mRNA and a new reinforcement learning based paradigm for mRNA sequence design.

## 1. Main text

Protein-coding mRNA sequences (codon sequences) are structured in non-overlapping triplets while non-coding RNA do not. Despite the central role that codon sequences play in the central dogma of molecular biology and in practical applications such as mRNA therapeutics^1^ and protein manufacturing^2^, there is still a lack of a strong foundational language modelling approach for codons. Recent advances in large language models have shown promise in modeling biological sequences^3^, including proteins^4-6^, non-coding RNA^7,8^, and DNA^9^; yet, a generative foundation model for coding mRNA doesn’t exist. Coding mRNA sequences have a unique latent structure, redundancy, and organism-specific bias embedded within codon usage^10-13^.

Here, we introduce codonGPT, a codon-native generative transformer language model. codonGPT is trained on a GPT2 architecture from scratch using protein coding nucleotide sequences, using a vocabulary of nucleotide triplets. During inference, we applied pre-softmax logit masking to constrain generation to only those codons that preserve the original protein sequence. We evaluated codonGPT by comparing its generated sequences to native codon usage patterns of House-Keeping genes (*HKG*)^14^ and an in-depth analysis of two genes of variable functions: Human Leukocyte Antigen A (*HLA-A*)^15^ and Actin Beta (*ACTB*)^16^. Our results for the first time show that the model can produce biologically plausible codon sequences while maintaining synonymous fidelity without requiring explicit downstream tuning.

For the first time, we then demonstrated the use of reinforcement learning on a biological generative language model like codonGPT for a very important downstream application of codon optimization. This is the first time reinforcement learning on such models is being used for this task. For most therapeutic and synthetic gene design tasks, we do not want a generic model that works fine for most proteins. Rather for such high value applications that rely on a single focused protein, we want a high performing model for a single protein that a user can customize and optimize as they desire. Usually expression, stability, or a mix of various biological parameters must be optimized under multiple biological constraints. Thus, to further align codonGPT generated sequences with expression-optimized and stability-optimized properties, we experimented with a fine-tuned version of codonGPT via reinforcement learning on a protein-specific codon optimization task using customized reward variables. This design allowed the model to adapt to sequence-specific trade-offs such as codon usage patterns, local GC content, and RNA structural constraints that may differ significantly between proteins. Using two proteins, HLA-A and ACTB, as examples, we showed that fine tuning codonGPT via reinforcement learning generates such optimized codon sequences.

## 2. Results

### 2.1. Biological Structure Emerges from Unsupervised Codon Level Training of codonGPT

We first evaluated whether codonGPT, trained in a fully unsupervised manner, could learn biologically meaningful representations of codon sequences. The model was trained as a next-token predictor at the codon level, with no explicit supervision regarding amino acid identity, gene structure, or expression. We analyzed both its predictive performance and the structure of its learned embedding space to assess the extent to which codonGPT internalized biological signals.

#### 2.1.1. Codon embeddings reflect synonymous codon relationships

We investigated whether codonGPT learns to group codons that encode the same amino acid by analyzing its internal representation space. After training, we extracted the 64 codon embeddings from the model’s input layer and visualized them using t-SNE **(Fig. 1a)**. The two-dimensional projection, clustered together by K-means clustering, revealed evidence of biological structure. Codons encoding the same amino acid tended to cluster in localized regions, with clearer separation observed for some synonymous groups than others. For example, glycine (abbreviated as G) codons (GGT, GGC, GGA, GGG), and glutamine (abbreviated as Q) codons (CAA, CAG) appeared close together in the t-SNE space, suggesting that codonGPT learns to represent synonymous codons with similar embeddings.

**Fig. 1:**
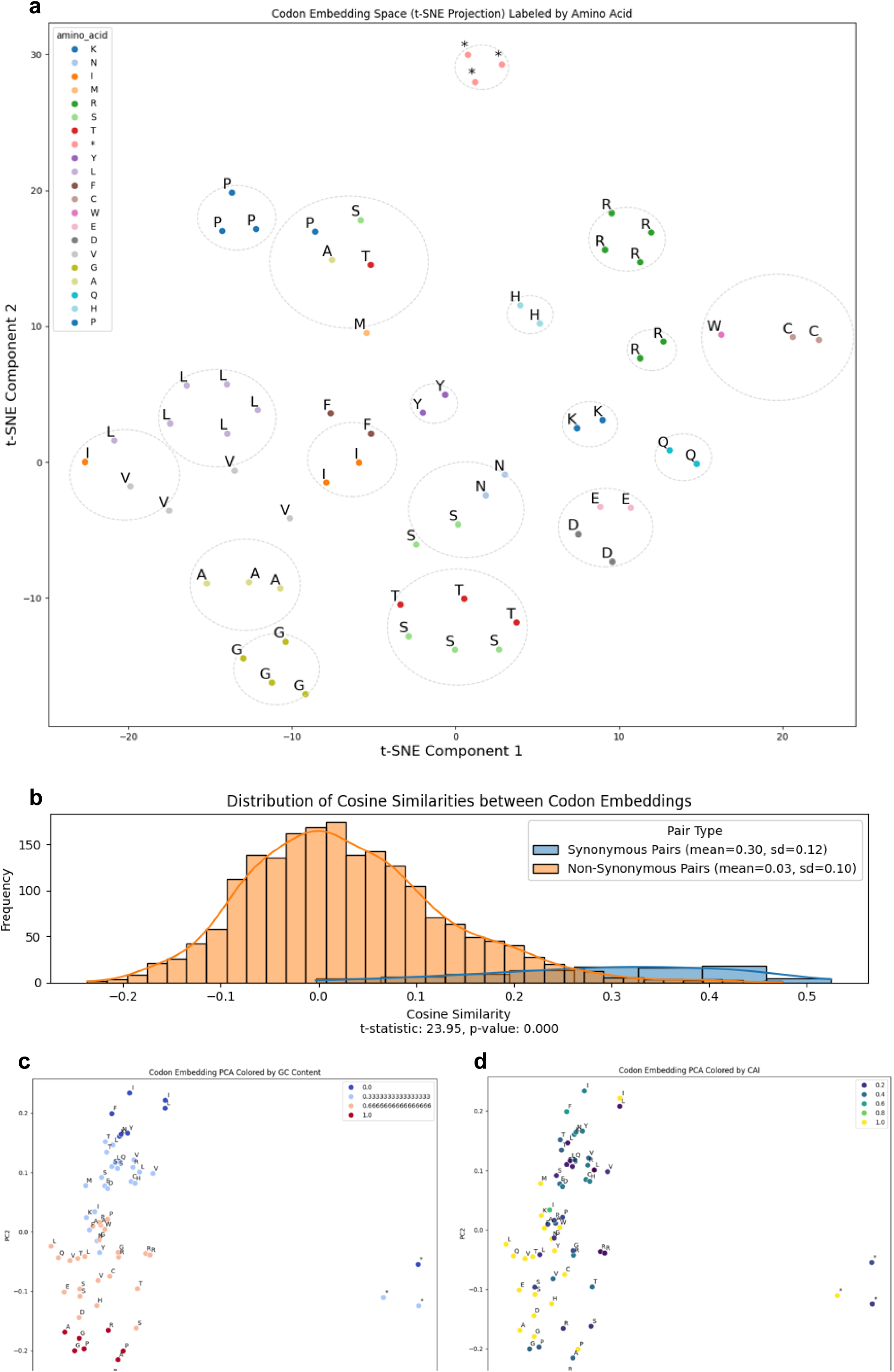
Analysis of codon embedding space of codonGPT. **a.** Codon embedding space projected onto two dimensions using t-SNE (n=2, perplexity = 10, random state = 42). Each point represents a codon, colored according to its corresponding amino acid. Circles highlight clusters identified by KMeans (n = 20 and random state = 42, demonstrating the model’s ability to group codons encoding the same amino acid. **b.** Cosine similarity distributions revealing the LLM’s learned representation of synonymous relationships. The histograms illustrate that synonymous codons tend to have higher cosine similarity scores, indicating more similar representations. The average cosine similarity for synonymous pairs (0.203±0.102), significantly higher (t = 19.238, p < 7.710e-76) than that for non-synonymous pairs (−0.010±0.101). **c.** PCA plot colored by GC content, highlighting the distribution of codons based on base composition, with low GC-content codons positioned higher and high GC-content codons positioned lower, reflecting genomic stability and codon context. **d.** PCA plot colored by Codon Adaptation Index (CAI), demonstrating a linear arrangement reflecting codon usage bias, with groupings of codons based on CAI values and translational efficiency. Notably, stop codons in **a, c, d** plots appear as outliers, indicating their distinct sequence and functional properties.

To determine whether this visual structure reflected a statistically meaningful pattern, we computed the average pairwise cosine similarity between codons within each synonymous group and compared it to the average similarity of non-synonymous codon pairs. The mean cosine similarity between synonymous codons was 0.30, while non-synonymous codon pairs had a near-zero average similarity (0.03) **(Fig. 1b)**. The resulting difference of +0.27 (p<0.05) demonstrates that synonymous codons are consistently embedded more closely together than non-synonymous codons. This pattern emerged solely from the model’s unsupervised training objective, without any explicit access to amino acid labels or translation supervision. The proximity of synonymous codons in the embedding space suggests that codonGPT internalizes semantic relationships embedded in the structure of protein-coding sequences. Rather than relying solely on surface-level codon frequencies, the model appears to develop a deeper representation of codon meaning based on the broader sequence context in which codons appear. Importantly, the synonymous relationships captured in the embedding space mirror the structure of the genetic code, despite the model having no direct knowledge of amino acid translation or protein function.

These results demonstrate that codonGPT learns biologically meaningful structure at the level of codon synonymy, and that this structure is reflected both qualitatively by tSNE and quantitatively by cosine similarity in its learned representation space. The emergent clustering of synonymous codons highlights the model’s potential as a foundation for biologically constrained generation, synonym-aware optimization, and tasks requiring semantic understanding of codon usage.

#### 2.1.2. Compositional biases encoded in embedding geometry

To determine whether codonGPT embeddings reflect lower-level nucleotide composition, particularly GC content, which plays a key role in genome structure and gene regulation. We colored the codon PCA space by GC percentage and observed a smooth gradient along one of the principal axes **(Fig. 1c)**. GC-rich codons clustered at one end of the space, while AT-rich codons were positioned at the opposite end. The continuity of this gradient suggests that compositional properties are embedded directly into the codon representations, likely due to the model’s sensitivity to local sequence features during training.

#### 2.1.3. Expression-related codon usage properties emerge from unsupervised training

We then investigated whether codonGPT representations align with codon usage preferences that are functionally relevant to gene expression. Using CAI values derived from highly expressed human genes, we colored the PCA projection by CAI score. Strikingly, codons with high CAI values formed a distinct, compact region in the embedding space, whereas lower-CAI codons appeared more dispersed **(Fig. 1d)**. This spatial organization implies that codonGPT learns codon preferences that align with translational efficiency—even though it has never seen expression data or codon usage labels.

Together, these results indicate that codonGPT, trained without any explicit biological supervision, learns codon-level representations that encode translational relationships, compositional properties, and expression-related usage biases. This embedding geometry provides a strong foundation for downstream applications in synonymous codon selection, constrained decoding, and biologically informed optimization.

### 2.2. Inference-Time Synonymous Logit Masking of codonGPT Ensures Biologically Faithful Codon Generation

We then enhanced our codon-level language model with inference-time, pre-softmax, synonymous logit masking. This approach constrains the model such that at each generation step, it selects only from codons that correspond to the correct amino acid, ensuring the biological meaning of the sequence is preserved. We evaluated this approach by comparing a set of native codon sequences of 100 protein coding house-keeping genes against those generated by our model for those same genes. Our evaluation focused on both fidelity to the target protein and the biological plausibility of the generated codon sequences.

#### 2.2.1. Protein Sequence Preservation

The model preserved the target protein sequences with 100% accuracy in all 100 cases. This confirms that the pre-softmax logit masking mechanism effectively enforces the amino acid constraint throughout the entire decoding process. Since the model is restricted to generating only synonymous codons for each position, it ensures that the decoded protein is identical to the intended sequence. This guaranteed preservation is critical for biological applications such as codon optimization, gene synthesis, and therapeutic mRNA design, where maintaining the protein output is non-negotiable. Unlike unconstrained generation, which can result in mistranslation errors, our masking strategy fully eliminates this risk at inference time.

#### 2.2.2. Gene-specific codon bias is preserved while introducing biologically plausible diversity

For each gene, we compare the generated codon sequence to its native counterpart using several sequence-level and codon-level similarity metrics **(Fig. 2)**.

**Fig. 2:**
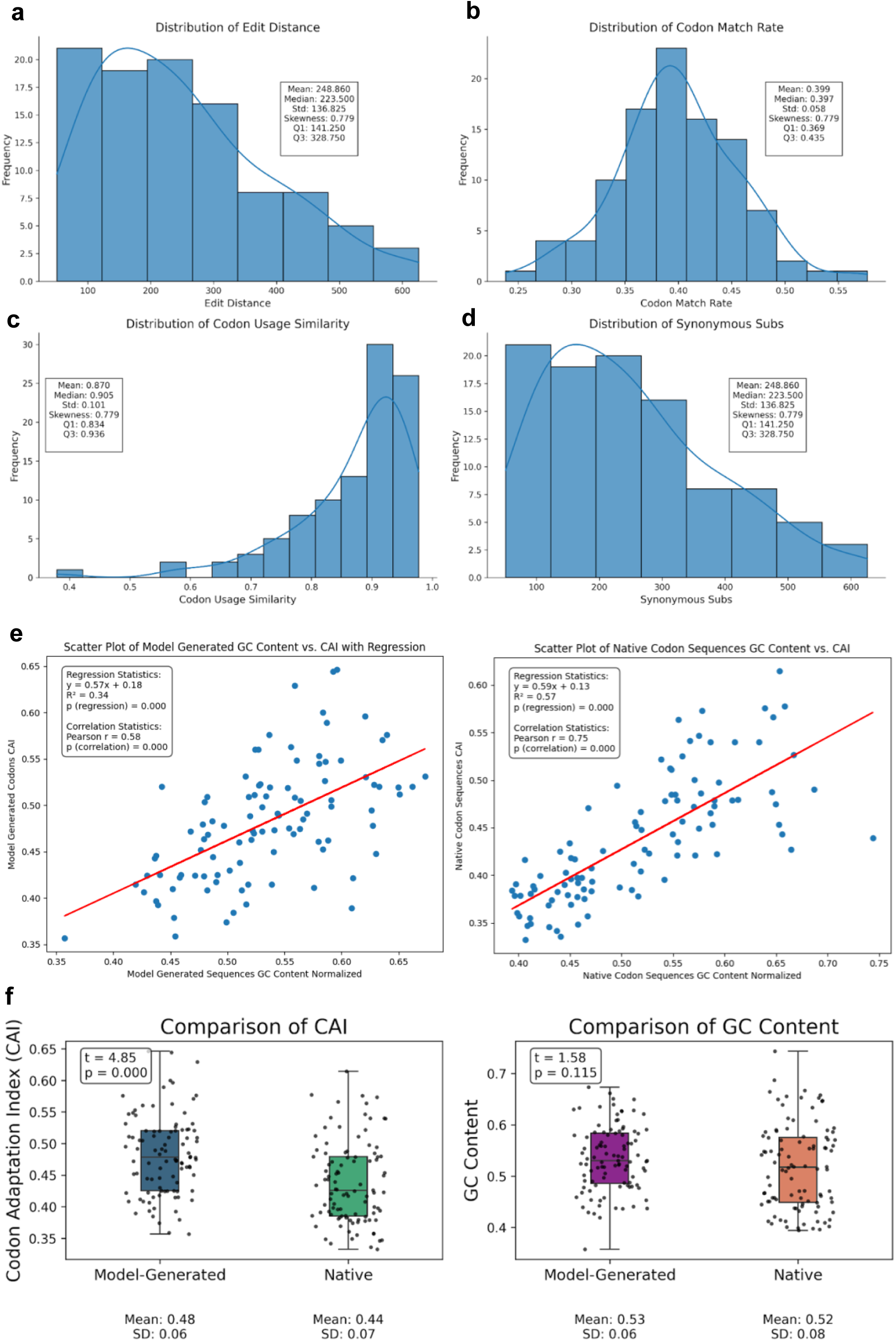
Analysis of codonGPT generated codon sequences of 100 unique house-keeping genes with native codon sequences of the same set of 100 unique house-keeping genes. **a.** Distribution of edit distances between native and generated codon sequences shows a median of 223 substitutions per gene. **b.** Codon match rate distributions reveal a median overlap of 39.7%, reflecting high sequence-level fidelity. **c.** Codon usage similarity, computed as cosine similarity between codon frequency vectors, is tightly clustered around native profiles (median = 0.87, IQR = 0.83–0.93), indicating the model’s ability to capture gene-specific codon bias. **d.** The number of synonymous substitutions per gene has a median of 223 (range: 141.25–328.75), confirming that the sequence diversity introduced by the model remains within biologically plausible limits while maintaining protein identity. **e.** GC content versus CAI shows a strong positive correlation in both model-generated (left) (Pearson’s r = 0.58, *p* < 1×10⁻⁶) and native(right) (Pearson’s r = 0.75, *p* < 1×10⁻⁶) sequences, indicating that the model preserves the native GC–CAI relationship. **f.** Box plots show slightly higher CAI content (left) between groups (mean = 0.48 vs. 0.44, t-test = 4.85, *p* < 1×10⁻⁶) while GC content (right) is not significantly different in model outputs (mean = 53% native vs. 52% generated, t-test = 1.58, *p* = 0.115).

The edit distance between native and generated sequences had a median of 223 substitutions per gene, highlighting that the model introduces substantial codon-level variation **(Fig. 2a)**. Importantly, this variation was largely synonymous, with the number of synonymous substitutions also centered around a median of 223 (range: 141.25– 328.75), ensuring the translated protein remains unchanged **(Fig. 2d)**.

Despite the sequence diversity, codon-level overlap remained moderate, with a median match rate of 39.7%, indicating partial conservation of specific codons **(Fig. 2b)**. More critically, the model preserved global codon usage trends per gene: cosine similarity between codon frequency vectors of native and generated sequences yielded a tightly distributed similarity profile (median = 0.87, IQR = 0.83–0.93) **(Fig. 2c)**, suggesting that codonGPT effectively learns gene-specific codon biases.

#### 2.2.3. Biological Quality of Generated Codon Sequences

We next examined whether codonGPT-generated sequences retain key biological correlations observed in native codon sequences of the house-keeping genes.

Specifically, we evaluated the relationship between GC content and CAI, two metrics known to be co-regulated in many human genes. Despite never explicitly optimizing for these metrics during training, the model produced codon sequences with highly realistic profiles. In both model-generated and native sequences, we observed a strong positive correlation between GC content and CAI (Pearson’s *r* = 0.58 and *r* = 0.75, respectively; *p* < 1×10⁻⁶ for both **(Fig. 2e, left and right)**, suggesting that the model internalizes and reproduces this underlying biological relationship.

To further quantify differences, we compared overall CAI and GC content between model generated and native sequences. While GC content was not significantly different between the two sets (mean = 53% model generated vs. 52% generated native; *t* = 1.58, *p* = 0.115) **(Fig. 2f, right box plot)**, CAI was modestly but significantly higher in the model generated sequences (mean = 0.48 vs. 0.44; *t* = 4.85, *p* < 1×10⁻⁶) **(Fig. 2f, left box plot)**. This indicates that while codonGPT preserves the native GC–CAI coupling, it may bias slightly toward higher translational efficiency. This suggests that the model not only respects fundamental biological constraints but may also subtly optimize codon usage patterns in a biologically meaningful direction.

These results are especially noteworthy because the model operates under hard constraints, it must choose from a subset of valid codons per amino acid, yet it still tends to favor codons that match human codon usage biases. This suggests that the model has implicitly learned organism-specific patterns of synonymous codon usage. Together, these results show that codonGPT balances codon-level novelty with biologically constrained fidelity—generating synonymous sequences that reflect endogenous usage patterns without altering protein translation.

### 2.3. Emergence of codon constraints across functionally distinct genes

To investigate whether the codonGPT internalizes biologically meaningful constraints on synonymous codon usage, we performed a focused analysis of model-generated codon sequences for two structurally and functionally distinct human genes: a major histocompatibility complex class I gene (*HLA-A*), and β-actin, a housekeeping gene (*ACTB*). These genes were selected to represent distinct evolutionary pressures: immune-domain modularity in *HLA-A*, and constitutive expression stability in *ACTB*. For each gene, we generated 100 codon candidates using synonym-constrained decoding, where the amino acid sequence is fixed and each codon position is sampled only from synonymous alternatives. These candidates were generated using the base model without top-*k* filtering and with temperature set to 1.5 (*HLA-A*) and 1.0 (*ACTB*).

We first examined positional entropy across codon sites to evaluate per-position codon variability. *ACTB* exhibited higher average positional entropy (mean = 1.43) compared to *HLA-A* (mean = 1.31), reflecting gene-specific differences in synonymous codon flexibility **(Fig. 3a)**. The distribution of entropy values showed moderate variability in both genes (SD = 0.64 for HLA-A, 0.65 for *ACTB*) without substantial skew, suggesting relatively uniform per-site diversity **(Fig. 3b)**. Pairwise Levenshtein distances between generated sequences revealed non-trivial sequence-level diversity (mean = 205.98, SD = 23.80 for *HLA-A*; mean = 237.76, SD = 33.69 for *ACTB* **(Fig. 3c)**, confirming that codonGPT generates distinct synonymous variants rather than minor variations around a single motif. We also explored RNA secondary structure stability (ΔG) and repeat codon frequencies to assess structural and translational constraints. No strong patterns were observed between these two features **(Fig. 3d)**, suggesting they are not explicitly learned or prioritized by the model in the unsupervised setting. Lastly, generated candidate sequences were well within biologically realistic bounds of ∼0.3 to ∼0.6 for CAI of both genes and ∼0.4 to ∼0.65 for GC of both genes **(Fig. 3e)**, further demonstrating that codonGPT maintains translational and compositional plausibility.

**Fig. 3:**
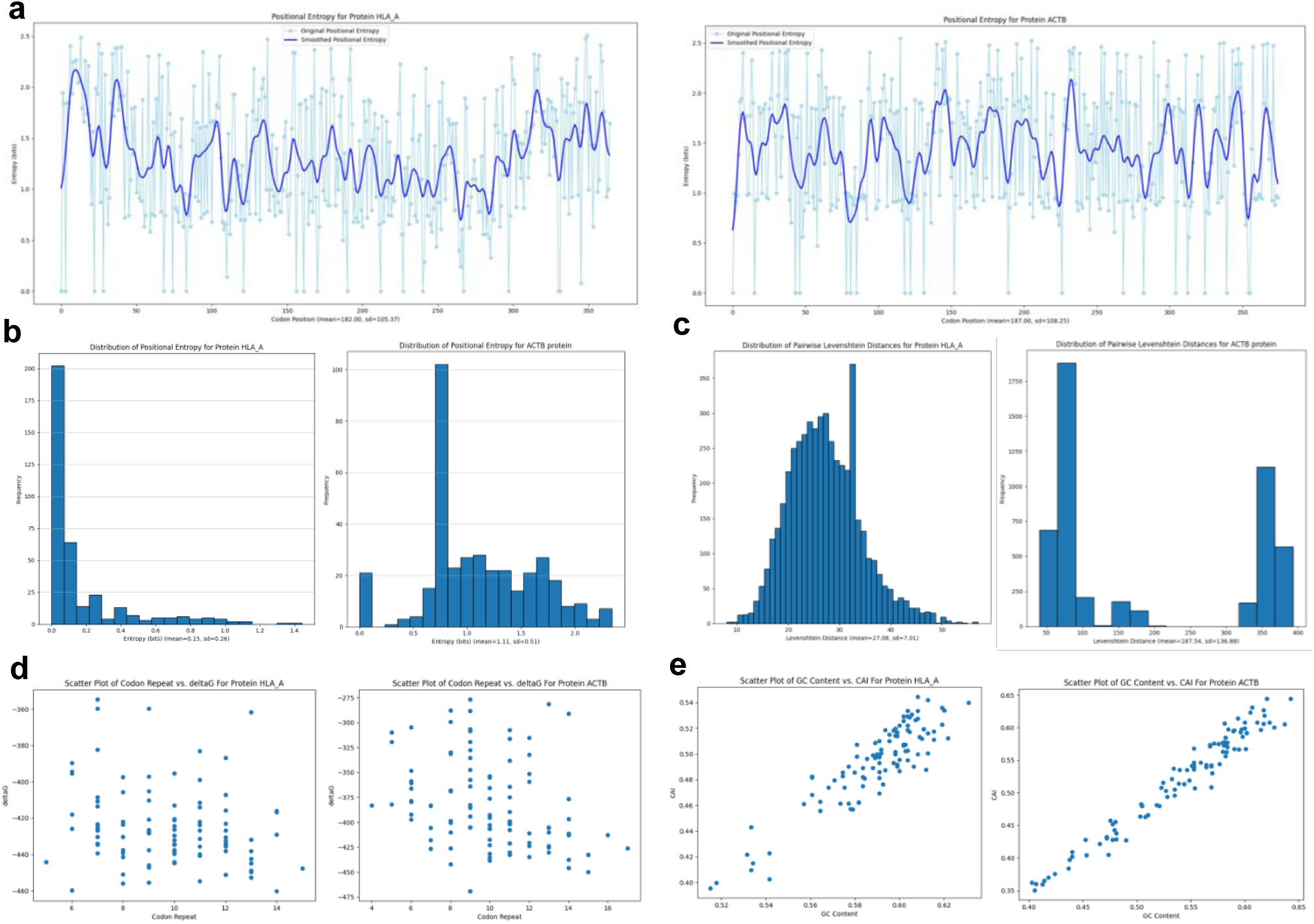
Analysis of 100 candidates generated from codonGPT for two genes: *HLA-A* and *ACTB*. **a.** Positional entropy for *HLA-A* (mean = 1.31) (left) and *ACTB* (mean = 1.43) (right) reflects gene-specific flexibility in synonymous codon usage, with *ACTB* showing broader per-site variability. **b.** Distributions of entropy values show moderate variability (*HLA-A*: SD = 0.64; *ACTB*: SD = 0.65), with no pronounced skew. **c.** Levenshtein distances between generated sequences confirm non-trivial synonymous diversity (*HLA-A*: mean = 205.98, SD = 23.80; *ACTB*: mean = 237.76, SD = 33.69). **d.** ΔG values versus repeat codon frequencies show no strong trends, suggesting these features are not emphasized by the unsupervised model. **e.** CAI values versus GC content fall within biologically realistic ranges. Together, these results illustrate that the model captures both gene-specific and position-specific codon constraints, even under unsupervised training and synonym-only decoding. Comparisons for HLA-A (generated using codonGPT hyperparameter temperature =1.5, topk=None) and ACTB (generated using codonGPT hyperparameter temperature =1, topk=None).

Together, these results indicate that codonGPT captures both gene-level codon usage trends and position-specific variability, even in the absence of supervised fine-tuning highlighting the model’s ability to internalize and reflect the biological grammar of coding sequences.

### 2.4. Protein specific Reinforcement Fine Tuning of codonGPT optimizes a multi objective reward function

To use codonGPT on a downstream task of codon optimization, we fine-tuned codonGPT using protein-specific RL feedback for two representative proteins - HLA-A and ACTB. We constructed a reward function that combines 5 different biological components: CAI, which reflects translational efficiency in the host organism; GC content, which impacts mRNA stability and synthesis; RNA folding stability (ΔG score), a proxy for the structural integrity of the transcript; codon entropy, which measures synonymous codon diversity; and the number of codon repeats. These metrics were calculated for each training step and for each candidate generated during inference.

To evaluate the training stability and convergence behavior of our reinforcement learning framework, we analyzed and corrected the weights of our rewards based on the reward trajectory plot **(Fig. 4)**. We observed that the raw reward increased rapidly in the initial steps, sometimes followed by a valley, which later the model was able to extricate out of and then finally plateauing, indicating convergence to a locally optimal sequence. To reduce noise and better visualize the convergence pattern, we applied a moving average smoothing window, which revealed that most improvements occurred within the first 350–700 steps, with diminishing returns thereafter. This consistent plateau across various runs confirms that the model was able to reliably optimize for the multi-objective reward.

**Fig. 4:**
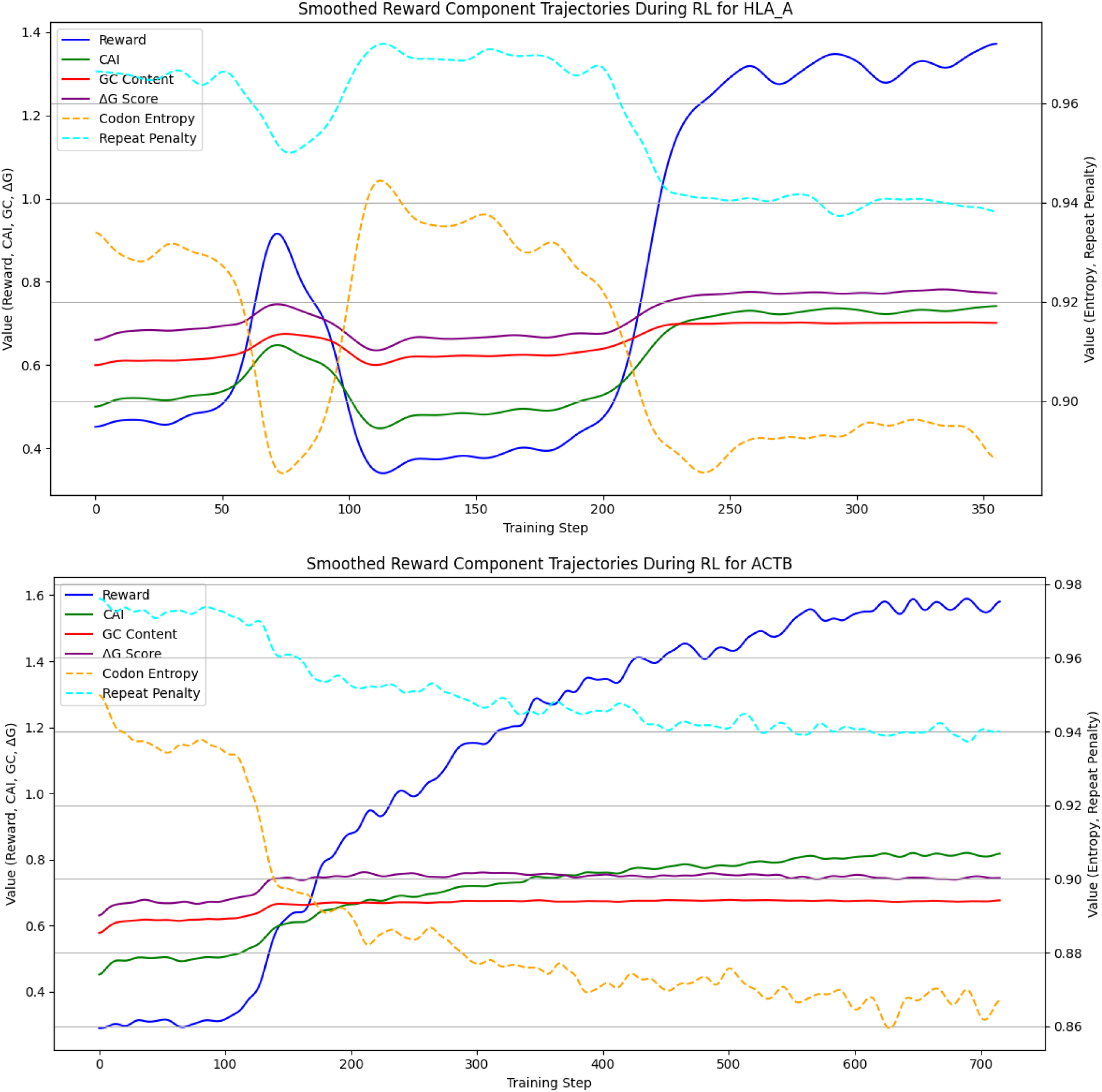
Training dynamics and outcomes of reinforcement learning–based codon optimization of two proteins: HLA-A, ACTB. Smoothed reward trajectory over training steps, illustrating convergence of both the total reward and its biologically grounded components, including codon adaptation index, GC content penalty, ΔG, codon entropy, and repeat penalty score. RL fine-tuning consistently improves codon sequence quality across all metrics, demonstrating biologically meaningful and stable optimization.

To evaluate the effectiveness of reinforcement learning (RL) for protein-specific codon optimization, we compared 100 candidate codon sequences generated by three models: a base codonGPT without any fine tuning, a codonGPT fine-tuned with protein-specific Reinforcement Learning (codonGPT RL-Optimized), and a BERT based Codon LLM by Fallahpour, et al^17^. We also used the native naturally occurring codon sequence for that protein as our baseline for these comparisons. All sequences were assessed using four biologically grounded reward metrics.

RL-optimized sequences for both proteins consistently improved overall reward scores **(Fig. 5a)**. The RL-optimized sequences exhibited the highest average CAI (∼0.79 for HLA-A and ∼0.84 for ACTB). These sequences also maintained a GC content centered within 25% of the desired value of 50%. Their ΔG scores were consistently higher (∼0.78 for HLA-A and ∼0.67 for ACTB) than any of the other models, suggesting that the RL fine-tuning process selected for more stable RNA secondary structures. Importantly, codon entropy remained high (∼0.92 for HLA-A and ∼0.90 for ACTB), implying that optimization did not collapse the diversity of synonymous codon usage.

**Fig. 5:**
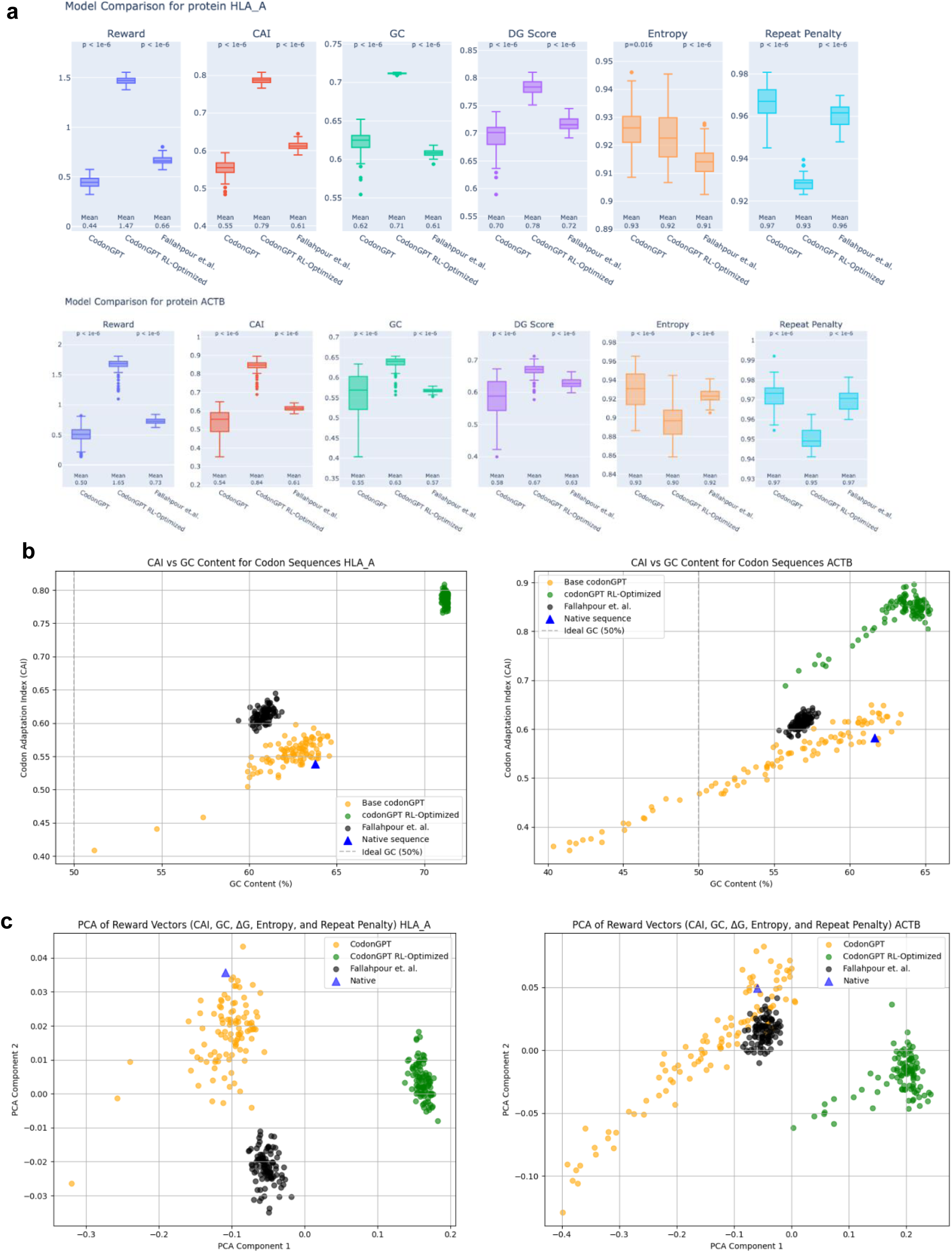
Structural and biological properties of RL-optimized codon sequences across two proteins: HLA-A, ACTB. **a.** Boxplots of reward components—CAI, GC content, ΔG, rare codon penalty, and codon repeat penalty—across sequences from the native gene, base codonGPT model, RL-optimized codonGPT model, and Fallahpour, et al’s method. For both proteins (HLA-A, top and ACTB, bottom panels, respectively), RL optimization of codonGPT increases the overall metric among all the other methods. For both proteins, it also balances each metric relative to baselines. Notably, RL sequences exhibit significantly higher CAI and higher stability (higher ΔG Score) than native or unsupervised outputs. **b.** Scatter plots of CAI versus GC content illustrate that RL-generated sequences cluster around high CAI values while maintaining GC content within 25% of ideal GC content. This balance suggests that the model learns to co-optimize translational efficiency and compositional stability. **c.** Principal Component Analysis (PCA) of reward vectors reveals that RL-generated sequences for both genes form a compact, high-reward cluster, distinct from the diffuse distributions of base codonGPT outputs. This shift reflects targeted optimization in multi-dimensional biological objective space. These effects are consistent across both genes and highlight the biologically grounded generalization capacity of the RL framework.

The interplay between translation efficiency and nucleotide composition was further examined by a scatter plot of CAI against GC content for all three models **(Fig. 5b)**. The RL-optimization was able to get the sequences to occupy a unique GC-CAI space not explored by the base codonGPT. This indicates successful resolution of a key biological trade-off between codon preference and synthesis feasibility.

We then evaluated how RL reshaped the generative distribution of the model by projecting the six-dimensional reward vectors into two principal components. The RL-optimized sequences formed a distinct, dense cluster within a high-reward region of the space, clearly separated from both the base model, the native sequence, or sequences optimized by Fallahpour, et al **(Fig. 5c)**. This concentration suggests that RL fine-tuning guides the model toward a narrow subspace of biologically optimized codon configurations—solutions that are unlikely to emerge from standard language modeling alone.

Collectively, these results demonstrate that reinforcement learning on a generative language model can produce codon sequences that are not only biologically interpretable and thermodynamically stable, but also compositionally balanced and robust across multiple biological objectives.

## 3. Discussion

The interface of RNA and proteins have been explored by recent deep learning approaches such as RNNs^18,19^, VAEs^20^, and GANs^21,22^. Several BERT approaches for codons^17,23,24^ have also recently emerged for codon sequence understanding. However, BERT’s bidirectional nature and masked pretraining objective^25^ make it less suited for tasks that require generative modeling. Usually in such techniques, a more refined dataset is used to fine tune the base model to teach it to optimize parameters the user wants, which is applied across all proteins. On the other hand, generative models, such as GPT, are able to generate full biologically valid sequences and have an inherent capability of being fine-tuned by reinforcement using any custom set of reward metrics a user might need to change as per protein. To address this gap in biologically aware codon design and enable flexible optimization of gene sequences, we presented codonGPT, a first of its kind, an unsupervised generative transformer model trained solely on human codon sequences.

This study is the first implementation of a generative architecture for codon level biological languages. Without alignment to protein sequences, the model captures codon usage bias, synonymous codon clustering, and long-range dependencies, demonstrating its ability to learn biologically meaningful patterns from raw sequence data.

To solve for amino acid fidelity during generation, we introduced a novel method of inference-time, pre-softmax, logit masking mechanism that restricts outputs to synonymous codons per position. This ensures semantic correctness without modifying model weights. We would like to point out that the customized logits processor based mechanism we proposed in this study also enables incorporation of additional biological constraints (e.g., restriction enzyme site detection, motif presence) at inference time itself without necessarily adding reinforcement learning.

We show for the first time that mRNA sequence design is made easier by our method. We showed this by extending codonGPT via RL for various biological optimization tasks. For protein-specific optimization, we developed a RL framework that combines CAI, GC balance, ΔG, and codon entropy into a unified reward function. This approach allows tailored optimization per protein, reflecting application needs in synthetic biology and therapeutic protein design. This user-customized balance between fidelity and variability is critical for biological design robustness. Fixed reward weights were used, but future work may explore adaptive strategies informed by empirical data. codonGPT can also be fine-tuned to additional design constraints not implemented in this study such as immunogenicity, ribosomal pausing, or tissue-specific translation.

Overall, codonGPT is the first of its kind to offer a scalable, modular, and biologically grounded framework for codon-level sequence generation and optimization. It serves as a foundation model for biologically constrained design, supporting both general-purpose biological discovery and target-specific engineering of protein-coding DNA. Unlike DNA- or protein-centric models, codonGPT provides a foundation model for sequences at the codon level and learns to generate synonymous codon chains while adhering to user-customized biological constraints. Our study establishes a foundation for broader applications in synthetic biology, multi-host expression tuning, mRNA vaccine design, and therapeutic antibody design.

## 5. Model availability

Our model is available at https://huggingface.co/anuj2054/codonGPT.

## 6. Acknowledgements

We would like to thank Dr. Richard S. Sutton from the University of Alberta for helpful discussion and suggestions related to the model.

